# The role of antifreeze glycopeptides (AFGP) and polyvinyl alcohol/polyglycerol (X/Z-1000) cocktails as ice modulators during partial freezing of rat livers

**DOI:** 10.1101/2021.08.04.455092

**Authors:** Shannon N. Tessier, Omar Haque, Casie A. Pendexter, Stephanie E.J. Cronin, Lindong Weng, Heidi Yeh, James F. Markmann, Michael J. Taylor, Gregory M. Fahy, Mehmet Toner, Korkut Uygun

## Abstract

The current liver organ shortage has pushed the field of liver transplantation to develop new methods to prolong the preservation time of livers from the current clinical standard of static cold storage. Our approach, termed partial freezing, aims to induce a thermodynamically stable frozen state at deeper storage temperatures (−10°C to −15°C) than can be achieved with supercooling, while simultaneously maintaining a sufficient unfrozen fraction to limit dehydration and ice damage. This research first demonstrated that partially frozen glycerol treated rat livers were functionally similar after thawing from either −10 or −15°C with respect to subnormothermic machine perfusion metrics and histology. Next, we assessed the effect of adding either of two ice modulators, antifreeze glycoprotein (AFGP) and a polyvinyl alcohol/polyglycerol combination (X/Z-1000), on the viability and structural integrity of partially frozen rat livers compared to glycerol-only control livers. Results showed that AFGP livers had high levels of ATP and the least edema but suffered from significant endothelial cell damage. X/Z-1000 livers had the highest levels of ATP and energy charge (EC) but also demonstrated endothelial damage and post-thaw edema. Glycerol-only control livers exhibited the least DNA damage on Terminal deoxynucleotidyl transferase dUTP nick end labeling (TUNEL) staining but also had the lowest levels of ATP and EC. Further research is necessary to optimize the ideal ice modulator cocktail for our partial-freezing protocol. Modifications to cryoprotective agent (CPA) combinations, as well as improvements to machine perfusion CPA loading and unloading, can help improve the viability of these partially frozen organs.

## I. Introduction

The liver organ shortage has pushed the field of transplantation to develop bold new strategies to preserve transplantable organs. Currently, the clinical standard of preserving transplantable livers is static cold storage (SCS) at 4°C, which keeps organs viable for a maximum of 12 hours [1]. Prolonging this preservation time would improve the allocation of organs in many ways. For example, it would reduce organ discard due to unacceptably long ischemic times, lower operating room costs by making liver transplant (LT) operations elective, enhance donor-recipient selection with human leukocyte antigen (HLA) matching and global matching programs, and make tolerance induction protocols more feasible [2–4].

Preservation methods to slow organ deterioration after procurement can be broadly categorized into two strategies: metabolic support and metabolic depression. Metabolic support through ex-vivo machine perfusion allows for continuous quality and viability assessment of organs. However, the major challenge with long term machine perfusion is maintaining organ homeostasis ex-vivo, which becomes exponentially more complex with longer perfusion durations that require continuous monitoring and adaptations [3,5–8]. On the other hand, metabolic depression strategies leverage the fact that tissue deterioration slows down at decreasing temperatures. Furthermore, lowering the hypothermic preservation temperature below 4°C harbors great potential to extend preservation times beyond clinical standards and does not require the long-term, constant maintenance of machine perfusion [9].

Within metabolic depression preservation strategies, most subzero preservation efforts have centered on low cryogenic temperature ranges (< −80°C) [3]. However, recent work has investigated prospects for expanding storage within the high subzero temperature range from −4°C to −20°C. These temperatures allow for more metabolic depression than hypothermic SCS while also potentially avoiding lethal ice formation and vitrification-related cryoprotectant toxicity and thermal stresses. The most prominent present example of the potential of this approach has involved storage below the thermodynamic freezing point at −4°C to −6°C in the absence of ice (termed supercooling), which has enabled 3-fold extensions of liver preservation duration [9–11]. While these studies have shown that ice formation in rodent and human livers can be completely circumvented with supercooling [9,11,12], the depth of metabolic stasis that can be achieved by this method is limited by the risk of spontaneous nucleation leading to damaging ice formation, which increases as temperature is lowered to below −6°C [13].

Although there is extensive evidence that ice formation can be severely damaging in tissues and organs [14–17], recent studies based on the survival of frozen animals such as frogs and turtles in nature have suggested that carefully limited and controlled ice formation may be tolerable during attempted cryopreservation of solid organs [2,18]. Ishine et al. showed that ice modulators such as antifreeze glycoproteins (AFGP) can have protective effects in high subzero liver freezing protocols by inhibiting ice recrystallization and preventing ionic leakage through cell membranes at low temperatures, although the authors report damage to the endothelial layer. The authors froze their rat livers for 2 hours used livers frozen with glycerol as the only cryoprotective agent (CPA) as controls [19]. To expand on these efforts, we aimed to test extended preservation durations (up to 5 days) in the presence of two ice modulators, antifreeze glycoprotein (AFGP) and a polyvinyl alcohol/polyglycerol combination (X/Z-1000), for their ability to confer freeze tolerance of rodent livers. The inclusion of these agents is called for in part because although the total quantity of ice present during long term storage at a fixed temperature is constant, the ice may still cause injury due to recrystallization that could be overcome by ice modulators.

AFGP has been shown to inhibit both ice recrystallization and ice growth below T_M_ (the thermodynamic freezing point). These glycopeptides inhibit ice growth by attaching to multiple faces of ice crystals [20–23]. AFGPs have also been shown to raise the homogenous ice nucleation temperature (T_H_) by organizing water into a more ice-like state [24]. However, since the temperature range in our partial freezing protocol is well above TH [25], the issues at hand involve the role of AFGP in ice shaping and ice recrystallization inhibition. Although AFGP can shape ice into damaging spicules [26], this effect may be outweighed under our storage conditions by the ice growth and recrystallization inhibitory effects of AFGP. X-1000 is a 2 kilodalton (kDa) polyvinyl alcohol [27] that contains 20% vinyl acetate, which improves the solubility and ice-inhibiting effects of X-1000 presumably by preventing self-association between X-1000 chains. Polyvinyl alcohol is known to inhibit ice recrystallization [28,29]. Z-1000 is a polyglycerol that inhibits heterogeneous ice nucleation [30], and together X/Z-1000 has been shown to protect rat hearts during supercooling [31–33] and is functional from 0°C to temperatures below −120°C [16].

In the present study, whole rat livers were frozen for up to 5 days at high subzero temperatures (−10°C to −15°C) by combining glycerol and ice nucleating agents (INAs) with the use of subnormothermic machine perfusion (SNMP) at 21°C. Further, two ice modulators, antifreeze glycoprotein (AFGP) and a polyvinyl alcohol/polyglycerol combination (X/Z-1000), were tested. Livers frozen with the inclusion of either AFGP or X/Z-1000 were compared to the control group (with glycerol as the main CPA) with primary outcomes being perfusion metrics, ATP, energy charge (EC), weight gain, and histology. We call this protocol “partial freezing” since it induces a thermodynamically stable frozen state at deeper storage temperatures (as low as −15°C) than can be achieved with supercooling, while simultaneously maintaining a sufficient unfrozen fraction to limit dehydration and ice damage.

## II. Materials and Methods

### A. Experimental design

**Figure 1** outlines the rat liver partial freezing protocol in 8 consecutive steps: (1) liver procurement, (2) preconditioning during SNMP, (3) CPA preloading during hypothermic machine perfusion (HMP), (4) loading of final storage solution and ice modulators during HMP, (5) partial freezing of rat liver, (6) thawing of rat liver, (7) unloading of CPAs during HMP, and (8) functional recovery of frozen rat livers during SNMP.

**Fig 1.**
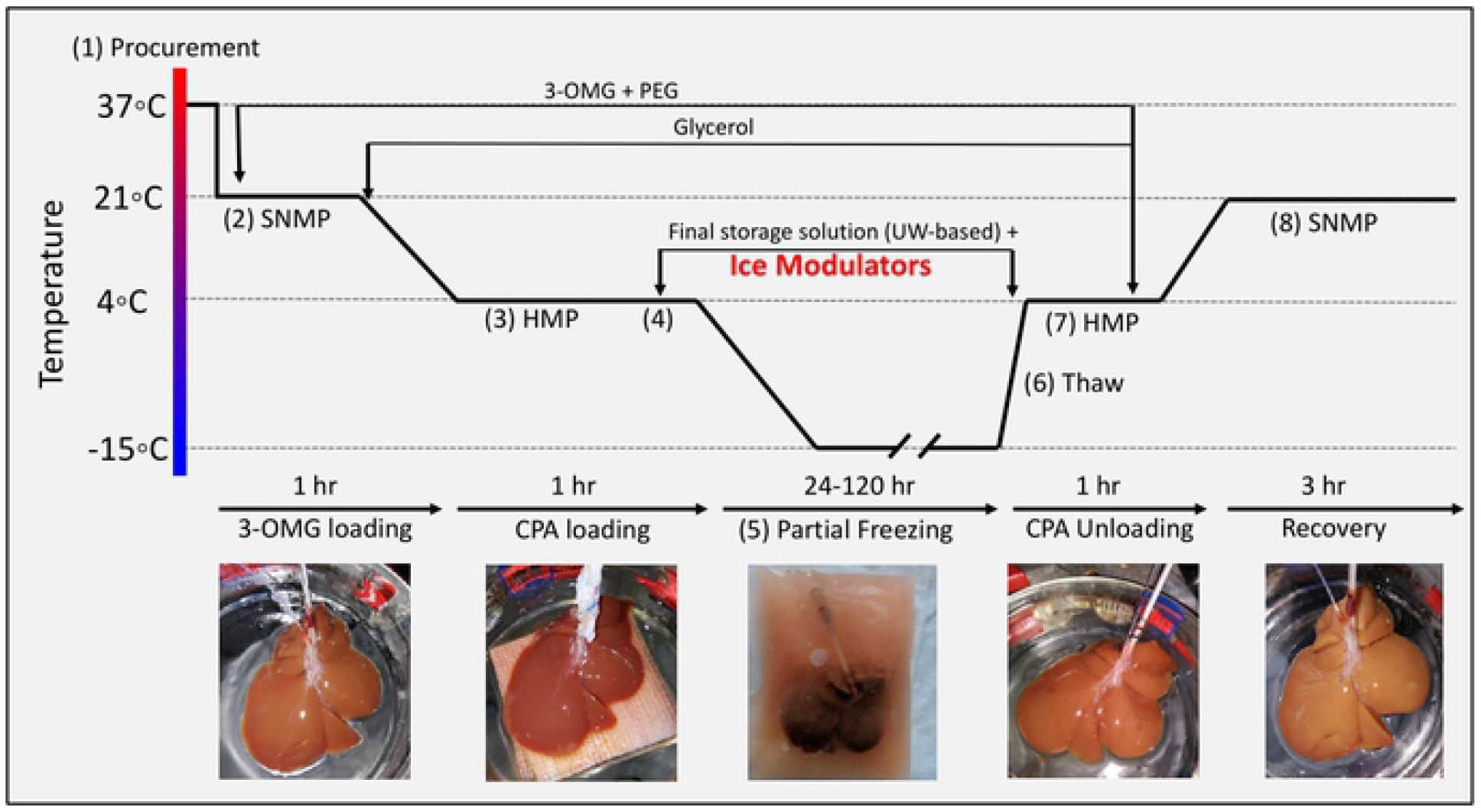
Experimental design of partial freezing, in 8 consecutive steps. (1) liver procurement, (2) preconditioning during SNMP, (3) CPA preloading during hypothermic machine perfusion (HMP), (4) loading of final storage solution and ice modulators during HMP, (5) partial freezing of rat liver, (6) thawing of rat liver, (7) unloading of CPAs during HMP, and (8) functional recovery of frozen rat livers during SNMP.

Within this protocol, we first compared partially frozen livers at −10°C (n=4 livers) and −15°C (n=9 livers) with 12% glycerol. Upon finding minimal differences between these two groups, we combined them as a control and compared them to livers partially frozen with 0.5 mg/ml (0.05% w/v) of AFGP (n = 4 livers) or 0.1% X-1000/0.2% Z-1000 (total, 0.3% w/v; n = 4 livers) ice modulating agents added to the preservation solution. After freezing, liver viability on SNMP was compared between the 12% glycerol control group and the two ice modulated groups.

### B. Partial freezing protocol

The rat liver perfusion system involved perfusion through the cannulated portal vein (PV) with regulation of pressure, flow, and temperature. Detailed set-up of the perfusion system has been previously described [34]. The total rat hepatectomy protocol was approved by the Institutional Animal Care and Use Committee (IACUC) of Massachusetts General Hospital (Boston, MA, USA). Livers were procured from male Lewis rats (250-300g, age 10-12 weeks. Charles River Laboratories, Wilmington, MA, USA) (**figure 1, step 1**). The bile duct was isolated and cannulated, and the rats were heparinized with 30U sodium heparin. PV splenic and gastric branches, as well as the hepatic artery were ligated prior to cannulation. The PV was subsequently cannulated with a 16-gauge catheter and immediately flushed with 40ml of heparinized saline (1000U/ml at room temperature). The liver was then freed from all peritoneal attachments, flushed with an additional 20ml of heparinized saline. After procurement, the liver was weighed, and machine perfusion was immediately initiated.

Preconditioning during SNMP was initiated at 21°C with a flow rate of 5 ml/min (**figure 1, step 2**). The flow rate was gradually increased [(1 ml/min)/min] until a maximum PV pressure of 5 mmHg, or a flow rate of 25 ml/min, was reached (whichever was reached first). The rat livers were perfused for 1 hour with 500 ml of preconditioning solution consisting of Williams E, 200 U/l of insulin, 2% w/v PEG, 50g/L BSA, 100mM 3-O-methylglucose (3-OMG), 10,000 U/l of heparin, 24 mg/L of dexamethasone, 25 mg/ml of hydrocortisone, 40,000 ug/l of penicillin, and 40,000 U/l of streptomycin and sodium bicarbonate as needed to maintain a physiological pH (see **Supplemental Table 1** for composition of all solutions).

After 1 hour of preconditioning during SNMP, the temperature was decreased to 4°C at a rate of ~1°C/min. Flow rates were gradually adjusted as necessary to ensure a maximum perfusion pressure of 3-5mmHg during HMP. The SNMP preconditioning solution was switched to 500 ml CPA preloading solution at 4°C (consisting of Williams E, 200 U/l of insulin, 2% PEG, 50g/L BSA, 100mM 3-OMG, 30mM raffinose, 3% hydroxyethyl starch (HES), 6% glycerol, 4000 U/l of heparin, 24 mg/l of dexamethasone, 25 mg/ml of hydrocortisone, 40,000 ug/l of penicillin, 40,000 U/l of streptomycin, and sodium bicarbonate as needed to maintain a physiological pH) (**figure 1, step 3; Table S1**). HMP was continued for 1 hour to ensure complete equilibration of solution throughout the liver parenchyma.

After CPA preloading during HMP, rat livers were loaded with 50mls of final storage solution (consisting of University of Wisconsin (UW) Solution (Bridge to Life Ltd., Columbia, SC, USA), 5% PEG, 50mM trehalose, 12% v/v (1.64M) glycerol, 1g/L Snomax (Telemet, Hunter, NY, USA) to promote nucleation, 10 U/l of insulin, 24 mg/l of dexamethasone, sodium bicarbonate as needed to maintain a physiological pH, and either 0.5 mg/ml of AFGP or 0.1% x-1000/0.2% z-1000 (**figure 1, step 4**). During the perfusion of the final storage solution the liver was perfused at a fixed flow rate of 0.5 ml/min for 30 min, resulting in a perfusion pressure of 3 mmHg.

Once the final storage solution during HMP had been perfused through the liver, the livers were placed in a storage bag with 50 ml of storage solution in a pre-cooled chiller (Engel, Schwertberg, Austria; model no. ENG65-B) for partial freezing (**figure 1, step 5**). Note: the concentrations of raffinose and HES were higher in the storage solution than in the pre-loading solution because the storage solution was made with UW which contains both raffinose and HES. In the case of livers frozen with glycerol only, the chiller temperature was pre-cooled to either −10°C or −15°C, and the liver was stored at one of these temperatures for 1 to 5 days. Livers frozen with either of the two ice modulation candidates were stored at −15°C.

After partial freezing, livers were thawed (**figure 1, step 6**) by placement in 50 ml of thawing solution consisting of Williams E, 10 U/l of insulin, 2% PEG, 50g/L BSA, 100mM 3-OMG, 30mM raffinose, 3% HES, 50mM trehalose, 5mM L-glutathione, 200μM cyclic AMP, 1,000 U/l of heparin, 24 mg/l of dexamethasone, 25 mg/ml of hydrocortisone, 40,000 ug/l of penicillin, and 40,000 U/l of streptomycin, which was pre-warmed to 37°C in a constant temperature bath (Thermo Fisher, Waltham, MA, USA). Livers were gently agitated until fully thawed, which required approximately 5 min based on superficial observation and thermal equilibration between the final thawing solution temperature and the liver surface temperature of 4°C.

After thawing, CPAs and INAs were unloaded during HMP (**figure 1, step 7**). The thawed livers were perfused at 4°C with thawing solution for 60 min with an initial flow rate of 2 ml/min. Flows were increased so as to keep a maximum pressure of 3mmHg. After 60 min, the perfusion temperature was increased to 21°C, and the perfusion solution was changed to 750 ml of SNMP recovery solution (consisting of Williams E, 20 U/l of insulin, 2% PEG, 50g/L BSA, 5mM L-glutathione, 200uM cyclic AMP, 1,000 U/l of heparin, 24 mg/l of dexamethasone, 25 mg/ml of hydrocortisone, 40000 ug/l of penicillin, and 40000 U/l of streptomycin; **Table S1**), discarding the first 50-100mls to ensure complete CPA removal. Functional recovery of frozen rat livers during SNMP with recovery solution continued for 3 hours (**figure 1, step 8**) with a maximum intrahepatic perfusion pressure of 5 mmHg and a maximum flow rate of 25 ml/min.

### C. Viability Assessment

Perfusate measurements were done hourly during the SNMP recovery period. Time zero (t=0) was defined as being at approximately 5 min of HMP, and the first outflow samples were taken at this time (flow, 2 ml/min). PV and infrahepatic vena cava (IVC) oxygen partial pressures and lactate levels were measured with a Cg4+ i-STAT cartridge (catalog no. 03P85-50) and handheld blood analyzer. Similarly, potassium and other electrolytes were measured in IVC samples using a Chem 8+ i-STAT cartridge (catalog no. 09P31-26) with the same blood analyzer (catalog no. WD7POC012; Abbott, Chicago, IL, USA).

Rat liver weight was measured directly after procurement, prior to freezing, after thawing, and after viability testing. Weight gain was calculated as the percentage increase at the end of recovery compared to the liver weight after procurement. Vascular resistance was calculated by dividing the perfusion pressure in the PV by the flow rate per gram of liver using the weight of the liver after procurement as the reference standard weight. Oxygen consumption rates were calculated as (pO_2_^in^ – pO_2_^out^)*F/W where pO_2_^in^ and pO_2_^out^ were the oxygen contents per ml of inflowing and outflowing perfusate, respectively, and the difference between them multiplied by the perfusion rate (F, in ml/min) provided the total oxygen uptake per minute. This value was then normalized by liver weight (W) to calculate the oxygen uptake per minute, per gram of liver. After thawing and SNMP recovery, rat liver tissue was either flash frozen in liquid nitrogen or fixed in 10% formalin, embedded in paraffin, sectioned, and stained with Hematoxylin and Eosin (H&E). Terminal deoxynucleotidyl transferase dUTP nick end labeling (TUNEL) was also performed on rat liver tissue after freezing to detect DNA breaks as an indicator of apoptosis [9]. Liver tissue that was flash frozen was used to quantify ATP and EC by the Massachusetts General Hospital (MGH) Mass Spectrometry Core (Boston, MA, USA).

Statistical analysis was performed with Prism 8 software (GraphPad Software, San Diego, CA, USA) with a significance level of 0.05. Analysis of variance (ANOVA), followed by Tukey’s post-hoc test (ANOVA/Tukey) was used for the comparison of the time-course perfusion data. ATP and EC in the −10°C vs. −15°C group were compared using unpaired, two-tailed t-tests.

## III. Results

### A. Comparison of partially frozen rat livers at −10°C vs −15°C with 12% glycerol

Pooled results for livers stored for 1 and 5 days at −10°C (n=4) or −15°C (n=9) with a cocktail containing 12% glycerol were compared 1 hour after CPA unloading and 3 hours after recovery with SNMP at 21°C. There was no statistically significant difference between the two groups with respect to oxygen consumption (**figure 2A**), PV lactate (**figure 2B**), vascular resistance (**figure 2C**), and perfusate potassium on two-way ANOVA/Tukey testing. Similarly, there were no statistically significant differences between −10°C and −15°C livers with respect to percent weight gain (**figure 2D**), ATP (**figure 2E**), and EC (**figure 2F**) by unpaired, two-tailed t-testing. Thus, rat livers partially frozen with 12% glycerol at −10°C and −15°C were not functionally different, and as a result, were grouped together to form a more statistically robust control group (n = 13) for comparison against the ice modulator groups.

**Fig 2.**
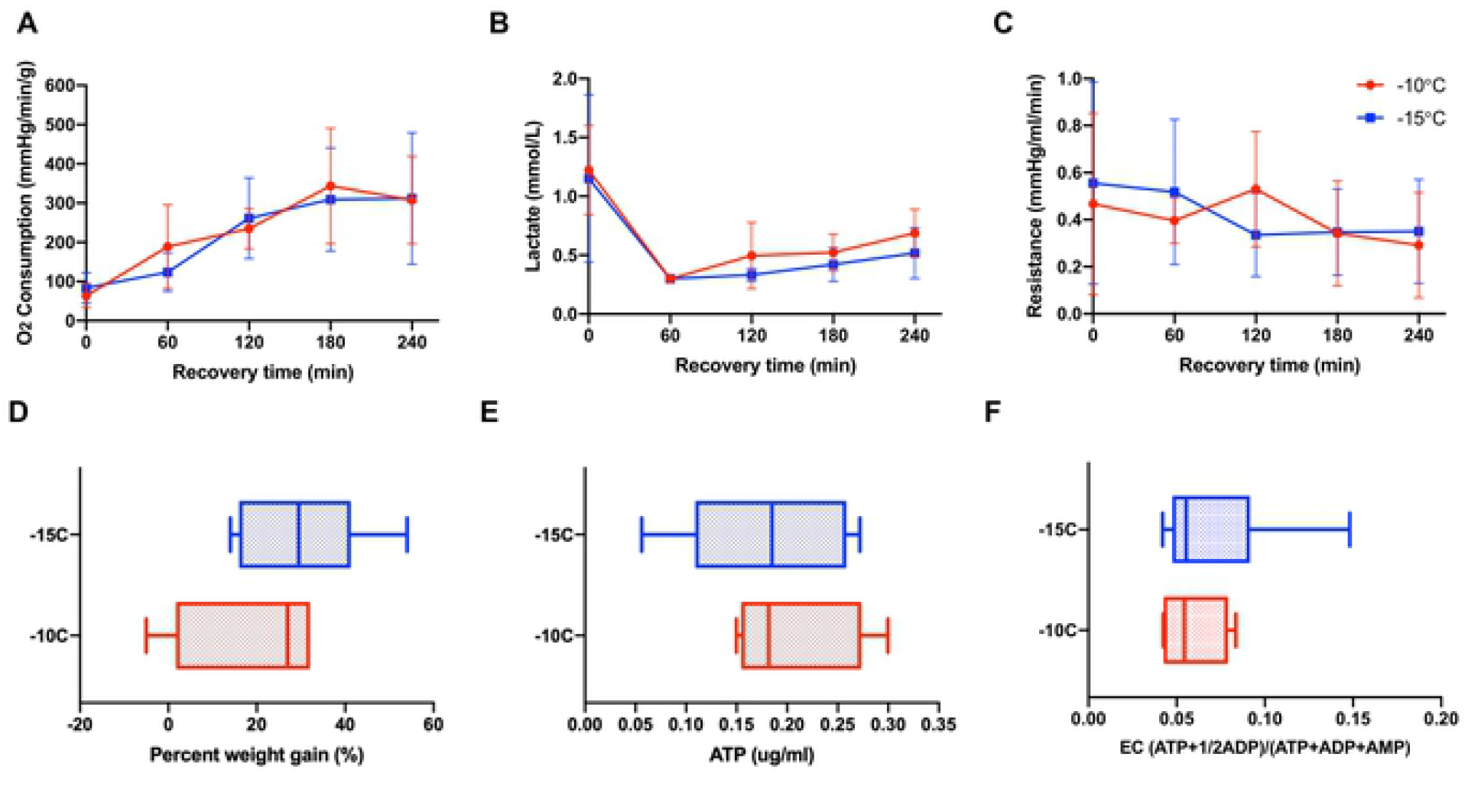
Perfusion metrics comparing partially frozen rat livers stored for 1 (n=6) and 5 (n=7) days at −10°C (n = 4) vs −15°C (n = 9) with 12% glycerol and 4 hours of recovery during SNMP revealed no functional differences between the groups. (A) oxygen consumption, defined as outflow oxygen minus inflow oxygen, adjusted for liver weight and flow rate, (B) inflow lactate, (C) resistance, defined as the perfusion pressure in the PV divided by the flow rate and adjusted for liver weight after procurement, (D) percent weight gain, defined as percentage increase of liver weight at the end of recovery compared to liver weight after procurement, (E) ATP, (F) energy charge. Two-way ANOVA, followed by Tukey’s post-hoc test for A-C. Unpaired two-tailed t-test for D-F. Boxes: median with interquartile range. Whiskers: min to max. Significance level: p < 0.05.

### B. Comparison of partially frozen control rat livers to livers frozen with AFGP or X/Z-1000 ice modulators (pooled results for 1 and 5 days)

There was no statistically significant difference between the three groups with respect to oxygen consumption based on two-way ANOVA/Tukey testing (**figure 3A**). Control glycerol-only livers had the lowest mean perfusate lactate (1.17 ± 0.61 mmol/L) at time zero. Over the first hour of SNMP, there was a decrease in lactate among all three groups. There were no significant differences between the groups at any time point according to the two-way ANOVA/Tukey test (**figure 3B)**. X/Z-1000 frozen livers had significantly higher mean vascular resistance (1.44 ± 0.37 mmHg/[(ml/min)/g] at 0 hours compared to both 12% glycerol control livers (0.528 ± 0.39 mmHg/ml/min, p-value 0.0315) and AFGP livers (0.413 ± 0.27, p-value 0.0215) (by ANOVA/Tukey). However, these differences were no longer significant at the remaining time points as the resistance levels converged over time (**figure 3C**). At 1 hour, glycerol only livers had higher mean perfusate potassium levels (6.5 ± 1.01 mmol/L) compared to both AFGP livers (5.1 ± 0.22 mmol/L, p-value 0.0167) and X/Z-1000 livers (5.6 ± 0.14 mmol/L, p-value 0.0275) (ANOVA/Tukey). However, these differences were no longer significant at the remaining time points as the potassium levels converged over time.

**Fig 3.**
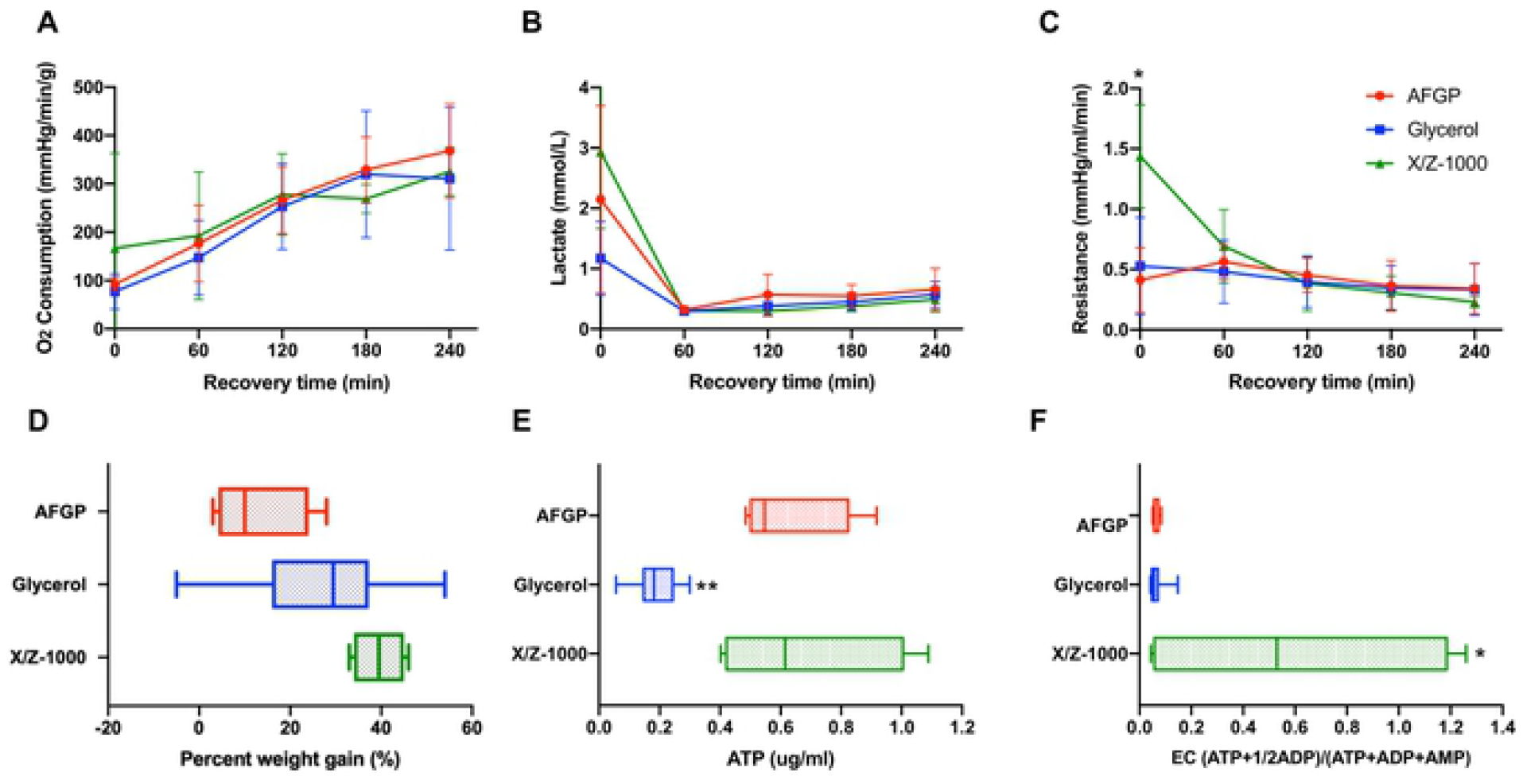
Perfusion metrics comparing rat livers partially frozen at −15°C with AFGP (n=4) or with X/Z-1000 ice modulators (n=4) versus 12% glycerol control (n=13), with 4 hours of recovery during SNMP. There was higher t=0 resistance and energy charge in X/Z-1000 livers, and lower ATP levels in glycerol livers. (A) oxygen consumption, defined as outflow oxygen minus inflow oxygen, adjusted for liver weight and flow rate, (B) portal vein lactate, (C) vascular resistance, defined as the perfusion pressure in the PV divided by the flow rate and adjusted for liver weight after procurement, (D) percent weight gain, defined as percentage increase of liver weight at the end of recovery compared to liver weight after procurement, (E) ATP, (F) energy charge. Stars denote statistical significance (two-way ANOVA, followed by Tukey’s post-hoc test for A-C, one-way ANOVA, followed by Tukey’s post-hoc test for D-F): *0.01 < p < 0.05; **0.001 < p < 0.01. Error bars represent standard deviation. Boxes: median with interquartile range. Whiskers: min to max.

Final mean weight gains for the glycerol-only control, the +AFGP, and the +X/Z-1000 groups were 26.9 ± 15.3%, 12.8 ± 13.2%, and 39.5 ± 5.69% respectively. Livers frozen with AFGP had the least edema, significantly less compared to X/Z-1000 livers (p-value 0.0294 by one-way ANOVA/Tukey; **figure 3D**). Mean ATP for the glycerol control group, AFGP, and X/Z-1000 were 0.187 ± 0.075, 0.624 ± 0.20, and 0.680 ± 0.32 ug/ml respectively. ATP concentrations were significantly lower for livers stored with glycerol only compared with AFGP (p = 0.0057) and X/Z-1000 (p = 0.0023) (ANOVA/Tukey). There was no significant difference in ATP levels between AFGP and X/Z-1000 frozen livers (p = 0.91) (**figure 3E).** Finally, mean EC for the glycerol control group, AFGP, and X/Z-1000 was 0.066 ± 0.032, 0.063 ± 0.015, and 0.591 ± 0.62 (ATP+1/2ADP)/(ATP+ADP+AMP) respectively. EC was dramatically higher in livers stored with X/Z-1000 and glycerol compared to glycerol alone (p = 0.0159) and to glycerol plus AFGP (p = 0.0428) (one-way ANOVA/Tukey). There was no significant difference in EC between AFGP and glycerol frozen livers (p = 0.99) (**figure 3F)**.

H&E staining of rat liver parenchyma following both 1 and 5 day partial freezing showed sinusoidal, hepatocellular, and endothelial cell damage in all groups. In glycerol plus X/Z-1000 frozen livers, H&E showed better preservation of sinusoidal patency (seemingly caused by less hepatocyte cell swelling) than seen in the other groups after 1 day of storage, which deteriorated somewhat after 5 days of storage. Endothelial patency also deteriorated between days 1 and 5, with almost complete endothelial cell destruction around the PV vasculature (after 1 day: **figure 4A**; after 5 days, **figure 4E**). After 1 day of storage, glycogen staining was decreased in glycerol plus X/Z-1000 livers, but not in the glycerol plus AFGP or glycerol-only groups. Glycerol plus AFGP frozen livers exhibited cellular swelling at the expense of sinusoidal patency and suffered from endothelial cell layer disruption that was mild after 1 day (**figure 4B**) and considerably worse after 5 days (**figure 4F**) of storage. Livers frozen with only glycerol at −10°C had sinusoidal damage and compression, peri-portal endothelial destruction (**figure 4D**) and linear cracks in the liver parenchyma (**figure 4H**). Decreasing the storage temperature of these control glycerol livers to −15°C did not alter the H&E pathology after either 1 (**figure 4C**) or 5 (**figure 4G**) days of frozen storage.

**Fig 4.**
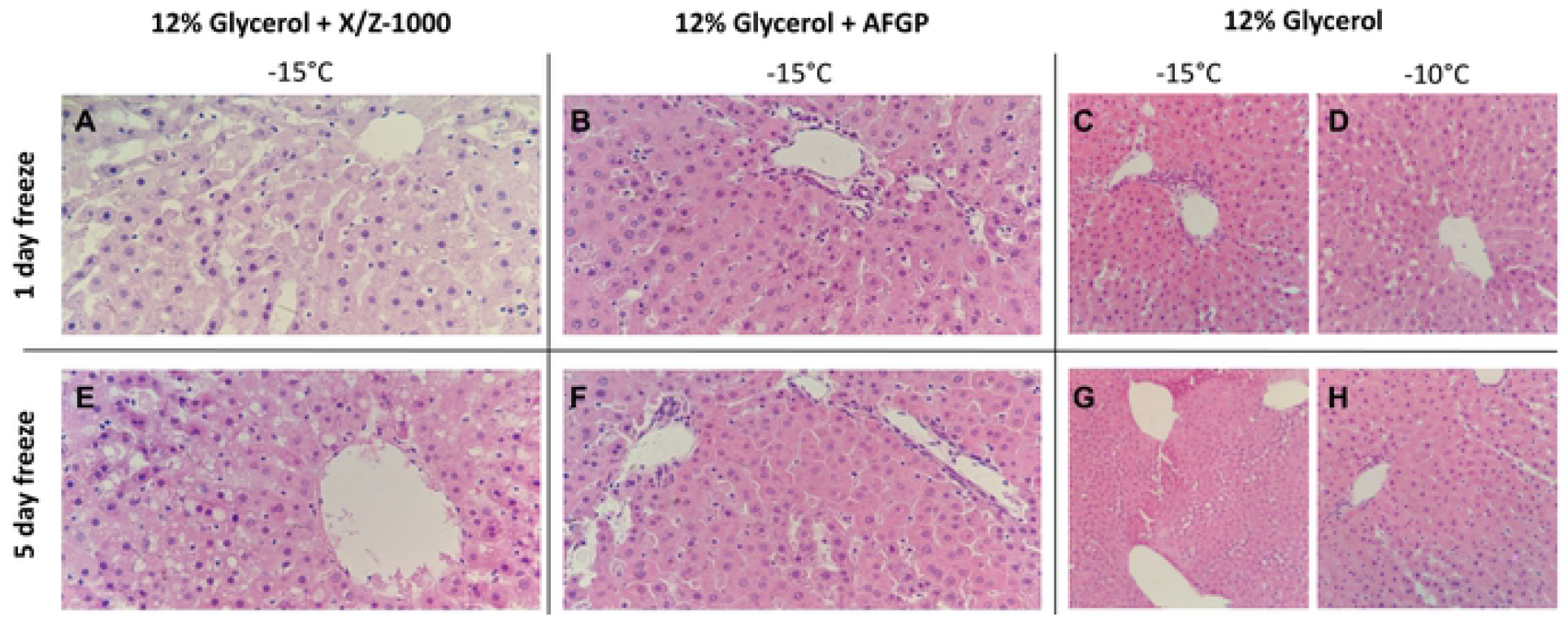
40X H&E staining of rat liver parenchyma after partial freezing for 1 day (A-D) and 5 days, (E-H) with X/Z-1000, AFGP ice modulators versus 12% glycerol only.

TUNEL staining was observed in both the endothelium and the sinusoids after 1 and 5 days of frozen storage in the presence of X/Z-1000, but appeared to be less intense after 5 days of storage (**figures 5A** and **5E**). AFGP frozen livers had primarily endothelial staining with mild sinusoidal staining after 1 day (**figure 5B**), but after 5 days, TUNEL staining had increased both around the vasculature and within the sinusoids (**figure 5F**). Compared to the other groups, AFGP frozen livers at −15°C and five days of partial freezing had the most TUNEL staining. Glycerol only livers stored at −10°C had remarkably less TUNEL staining compared to X/Z-1000 and AFGP livers with only mild sinusoidal staining after 1 day (**figure 5D**), and 5 days (**figure 5H**) of storage. As in the H&E results, decreasing the storage temperature of glycerol only livers to −15°C did not exacerbate TUNEL staining after 1 day (**figure 5C**) or 5 days (**figure 5G**) of storage.

**Fig 5.**
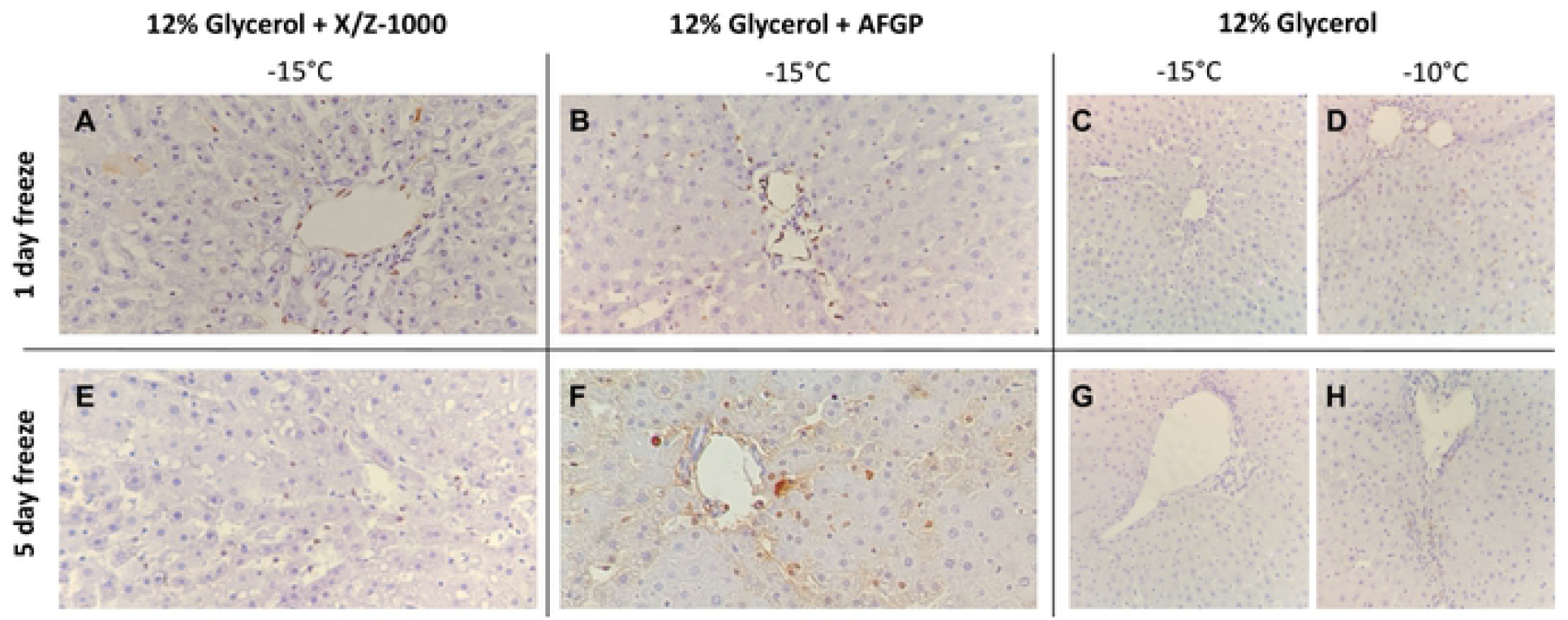
40X Terminal deoxynucleotidyl transferase dUTP nick end labeling (TUNEL) staining of rat liver parenchyma after partial freezing for 1 day (A-D) and 5 days (E-H) with X/Z-1000, AFGP ice modulators versus 12% glycerol only.

## IV. Discussion

Extending the preservation time of donor organs has tremendous clinical application in the field of transplantation. For liver transplantation, lengthening the allograft preservation time from the current standard of SCS at 4°C will reduce the burden placed on the healthcare system from high costs of unplanned surgeries, decrease organ rejection rates by incorporating more HLA typing into clinical practice, and possibly even open avenues for global matching programs [3,35].

Prior cryogenic organ preservation efforts have typically (although not universally [36]) encountered either lethal [14,15] or unacceptably damaging [16,37–39] amounts of extracellular ice, or the challenge of introducing the enormous concentrations of CPA needed to preclude such damage [16,40,41]. So far, these difficult challenges have not been adequately overcome, and therefore, other approaches should be investigated and may produce practical results more rapidly. Our current approach primarily utilized cold but not cryogenic temperatures to extend organ preservation time. We explored rat liver preservation at high subzero temperatures (−10°C and −15°C) combined with recovery SNMP to maximize the benefits of metabolic rate depression and *ex vivo* organ assessment, while avoiding the dangers of deep cryogenic temperatures. Ice modulators have been shown to modify ice crystal shape and inhibit ice recrystallization, potentially decreasing ice-induced cellular damage. In the context of the partial freezing of rat livers, ice modulators create an intriguing opportunity to preserve organs at subzero temperatures in the presence of ice, allowing these stored organs to reap the benefits of deeper metabolic stasis than current hypothermic standards while avoiding ice-related cellular damage.

This study first demonstrated that rat livers frozen at −10°C versus −15°C with glycerol were functionally similar regarding perfusion metrics, cellular architecture, and DNA damage, indicating that the reduction in partial freezing did not significantly reduce metabolic function. However, the reduction in storage temperature from −10°C to −15°C can have implications for organ viability on a cellular level. As water freezes, solutes are excluded from the ice crystals, which increases the osmolality and reduces the freezing point of the unfrozen water fraction. Thus, a lower freezing temperature results in more ice and a higher level of osmotic shift [42]. In our case, 12% v/v glycerol (1.64M), equates to 1.86 molality. According to the freezing point depression approximation, freezing point is lowered by about 1.86°C for every 1 osmolal increase in concentration. Adding the 0.3 osmolal contribution of the glycerol vehicle solution, the melting point of our storage media should be in the vicinity of −4°C. At −10°C, about 60% of the water in the solution will be converted to ice, and at −15°C, about 73% of the water will be frozen out, which is a significant increase. Although the membrane stabilizing saccharides, trehalose and raffinose [43] were employed, they do not enter cells and therefore do not nominally protect the inner membrane leaflet or reduce cell shrinkage induced by water extraction during freezing.

A major aim of this research was to assess the effect of ice modulators such as AFGP and X/Z-1000 on partially frozen rat livers compared to glycerol-only controls. AFGP frozen livers had the least amount of edema and high levels of ATP. However, the AFGP-mediated ice modulation had adverse effects on endothelial cells, which was reflected in both H&E and TUNEL staining, particularly after prolonged storage. As the duration of freezing increased from 1 to 5 days, the AFGP TUNEL staining expanded from predominately endothelial damage to diffuse sinusoidal cellular damage as well. AFGP has an established role in dynamic ice shaping, ice recrystallization inhibition, and hysteretic freezing point depression [22,44,45]. Since endothelial cells would make direct contact with the ice, it is possible that AFGP may be causing less favorable ice crystal shapes that are disrupting the endothelial cells of the liver. In a study by Rubinsky et al., antifreeze proteins resulted in the killing of all red blood cells during freezing despite the use of directional solidification methods that normally minimize ice damage [26,46]. Yet, AFGP may offer protection to hepatocytes through its other mechanism of action, ice recrystallization inhibition. (AFGP freezing point depression is typically limited to 1-2°C, which is smaller than the difference between our solutions’ T_M_s and our chosen storage temperatures and therefore was not able to contribute a protective effect in these experiments). Isothermal freeze fixation [15] could be useful in future studies for relating the details of ice distribution and characteristics to observed outcomes [47].

X/Z-1000 was the second ice modulator combination assessed in this study. X/Z-1000 frozen livers had the highest ATP and by far the highest EC but suffered from the highest SNMP resistance at t=0 and had the most edema after recovery. On staining, X/Z-1000 frozen livers had less TUNEL staining compared to AFGP frozen livers, but still exhibited both endothelial and sinusoidal staining in excess of that seen for the glycerol only group. X/Z-1000 livers had a large variation in both ATP and EC levels, which could potentially be explained by the competing mechanisms of action of X-1000/Z-1000 with the potent ice nucleator, Snomax [30]. Snomax would tend to reduce the number of ice crystals and, therefore, to increase their mean size and the range of grain sizes. This might relate to the larger observed size of the sinusoids and to stochastic differences in local nucleation and ice crystal size that affected the consistency of hepatocyte viability. While X/Z-1000 is appealing for its high ATP and EC, the high level of edema after partial freezing is concerning and might also be related to vascular damage caused by larger local intravascular or interstitial ice grains, which would be consistent with previous observations by Rubinsky et al. relating injury in frozen livers to vascular distension by intravascular ice [48]. A future direction related to this ice modulator should be to explore the use of X-1000, which is a recrystallization inhibitor, without the use of Z-1000, which is an antinucleator. On the other hand, total vascular distension should have been similar in all groups, as dictated by the phase diagram of glycerol-water solutions, and yet edema was more moderate in the glycerol-only group. In any case, X/Z-1000 seems promising for use with the isolation or preservation of isolated hepatocytes, for which maintenance of high ATP/EC would be the main goal.

Finally, glycerol-only control livers had the lowest lactate levels at t=0 and minimal TUNEL staining. However, these livers also had very low ATP and EC compared to the ice modulator groups. A potential biological reason for this difference is that glycerol induces glycerol kinase to convert glycerol to glycerol-3-phospahte, which is an ATP dependent pathway. Thus, the activity of glycerol as a CPA could depress ATP levels [49]. The extraordinary and unexpected ability of both ice modulators to prevent this has no clear explanation, but it seemed that utilization of glycogen to generate ATP and lactate was more effective in the X/Z-1000 group and to a lesser extent in the AFGP group, based on more intense glycogen staining (suggesting less glycogen metabolism) in the glycerol-only group. It would be interesting to see if structurally unrelated small molecule ice recrystallization inhibitors (IRIs) [50] would have a similar effect and also better protect the vascular system. Without the effects of the ice modulators preventing damaging ice recrystallization, glycerol-only livers also consistently had parenchymal cracks and endothelial cell obliteration on H&E staining, despite adding Snomax and 3-OMG to rescue endothelial cells from partial freezing injury. Thus, the concept of adding ice modulation to the basic methodology for high temperature freezing appears to be well supported. Finally, all liver groups cleared lactate over the 2 hour perfusion and (while incurring hepatocellular damage) had viable H&E histology after perfusion, meeting two criteria for transplantation [51,52]. Thus, future experiments transplanting these partially frozen livers after SNMP can be conducted to assess if perfusion performance corelates with *in vivo* hepatic function.

Overall, the ideal ice modulator combination to enhance the partial freezing protocol would retain the positive effects of high ATP and high EC seen in X/Z-1000, the low levels of edema with AFGP, without the cellular damage to endothelial and sinusoidal cells seen with both ice modulator groups. Thus, future directions to expand the preservation of livers for transplantation with the partial freezing approach depend on both modifications to the freezing protocol as well as the ice modulator combination. Specifically, increasing the CPA concentration to decrease ice formation at lower temperatures and improving the loading and unloading of CPAs with SNMP could improve liver viability. Additionally, there are other permeating CPAs that could be tested in the partial freezing protocol such as dimethyl sulfoxide, ethylene glycol, N-methylformamide, propylene glycol, and urea [40,53,54]. Finally, altering the base solution from UW to a lower potassium carrying solution could decrease the transmembrane osmotic stress in the unfrozen water fraction.

In summary, this research incorporated ice modulators into the rat liver partial freezing protocol to prolong the preservation time of livers. We demonstrated that there was no difference in partially frozen livers with only glycerol at −10°C versus −15°C with respect to perfusion metrics, cellular architecture, and DNA damage. Additionally, we showed that AFGP and X/Z-1000 ice modulators can have beneficial effects on partially frozen rat liver ATP and EC levels, respectively. However, further work on elucidating the optimal ice modulator cocktail is necessary as AFGP livers had high levels of endothelial DNA damage and X/Z-1000 livers suffered from post-freeze edema. Modifications to CPA combinations, as well as improvements to machine perfusion CPA loading and unloading, can help improve the viability of these partially frozen organs.

## Abbreviations

3-OMG: 3-O-methyl-D-glucose
ANOVA: Analysis of variance
AFGP: Antifreeze glycoprotein
BSA: Bovine serum albumin
CPA: Cryoprotective agent
EC: Energy charge
H&E: Hematoxylin and Eosin
HES: Hydroxyethyl starch,
HLA: Human leukocyte antigen
HMP: Hypothermic machine perfusion
INAs: Ice nucleating agents
IRIs: Ice recrystallization inhibitors
IVC: Infrahepatic vena cava
IACUC: Institutional Animal Care and Use Committee
LT: Liver transplant
M: molarity
PEG: Polyethylene glycol
X/Z-1000: Polyvinyl alcohol/polyglycerol
PV: Poral Vein
SCS: Static cold storage
SNMP: Subnormothermic machine perfusion
TUNEL: Terminal deoxynucleotidyl transferase dUTP nick end labeling
UW: University of Wisconsin
WE: Williams E

## Acknowledgements

This research was funded from the US National Institutes of Health (R01DK114506, R01DK096075, R01DK107875) and NSF ATP-Bio ERC grant (NSF 1941543). Further, we gratefully acknowledge funding to SNT for Career Development from NIH (K99HL1431149-01A1), American Heart Association (18CDA34110049), Harvard Medical School Eleanor and Miles Shore Fellowship, and the Claflin Distinguished Scholar Award on behalf of the MGH Executive Committee on Research. We gratefully acknowledge research support to Omar Haque by the American Liver Foundation (2019 Hans Popper Memorial Postdoctoral Research Fellowship) and the American College of Surgeons (Grant number 1123-39991 scholarship endowment fund).

## Data availability

The authors declare that the data supporting the findings of this study are available within the paper and its supplementary information files. Any additional data, if needed, will be provided upon request.

---

**Supplemental Table 1:** Composition of all solutions used in each phase of the partial freezing protocol. WE = Williams E, UW = University of Wisconsin, PEG = polyethylene glycol, BSA = bovine serum albumin, HES = hydroxyethyl starch, 3-OMG = 3-O-methyl-D-glucose, CPA = cryoprotective agent, INA = ice nucleating agents, U/l = Units per liter, *present in UW solution.

|  | Subnormothermic<br>Preconditioning<br>solution | Hypothermic<br>Preloading<br>solution | Storage<br>Solution | Thawing<br>Solution | Subnormothermic<br>Recovery<br>Solution |
| --- | --- | --- | --- | --- | --- |
| **Base solution** | **WE** | **WE** | **UW** | **WE** | **WE** |
| **Total volume** | **250 ml** | **250 ml** | **100 ml** | **250 ml** | **500 ml** |
| **Additives** |  |  |  |  |  |
| Insulin | 200 U/l | 200 U/l | 10 U/l | 10 U/l | 20 U/l |
| Heparin | 10,000 U/l | 4,000 U/l | --- | 1,000 U/l | 1,000 U/l |
| Dexamethasone | 24 mg/l | 24 mg/l | 24 mg/l | 24 mg/l | 24 mg/l |
| Hydrocortisone | 25 mg/ml | 25 mg/ml | --- | 25 mg/ml | 25 mg/ml |
| Penicillin | 40,000 ug/l | 40,000 ug/l | --- | 40,000 ug/l | 40,000 ug/l |
| Streptomycin | 40,000 U/l | 40,000 U/l | --- | 40,000 U/l | 40,000 U/l |
| Glutathione | --- | --- | 0.922 g/l* | 1.536 g/l | 1.536 g/l |
| **Macromolecules** |  |  |  |  |  |
| 35 kDa PEG | 20 g/l | 20 g/l | 50 g/l | 20 g/l | 20 g/l |
| BSA | 50 g/l | 50 g/l | --- | 50 g/l | 50 g/l |
| HES | --- | 30 g/l | 50 g/l* | 30 g/l | --- |
| **Saccharides** |  |  |  |  |  |
| 3-OMG | 19.42 g/l | 19.42 g/l | --- | 19.42 g/l | --- |
| Raffinose | --- | 15.12 g/l | 17.83 g/l* | 15.12 g/l | --- |
| Trehalose | --- | --- | 18.92 g/l | 18.92 g/l | --- |
| **CPA / INA** |  |  |  |  |  |
| Glycerol | --- | 60 ml/l | 120 ml/l | --- | --- |
| Snomax | --- | --- | 1 g/l | --- | --- |

**Supplemental Table 2:**
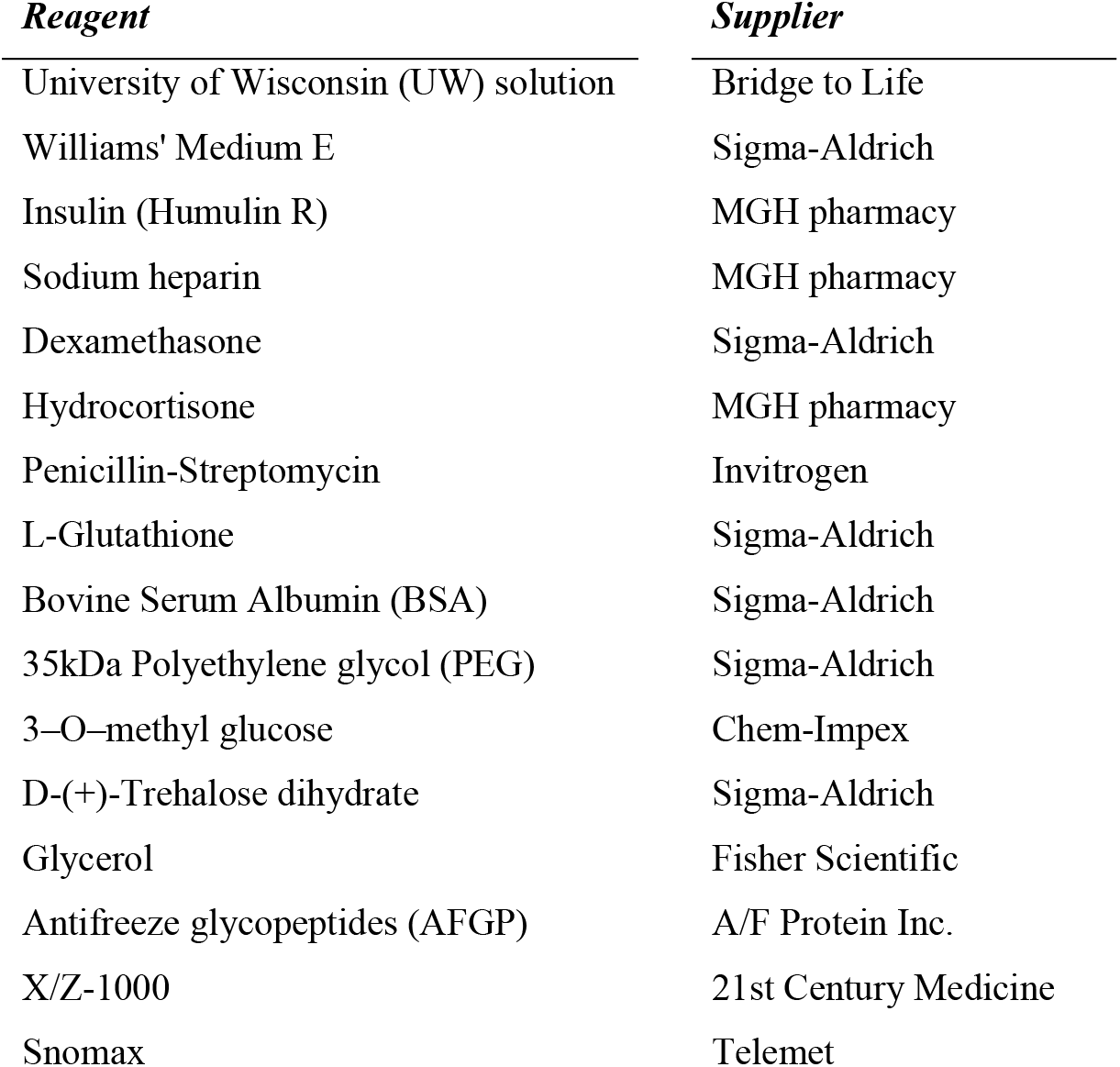
List of suppliers for each reagent used in partial freezing solutions.

| *Reagent* | *Supplier* |
| --- | --- |
| University of Wisconsin (UW) solution | Bridge to Life |
| Williams' Medium E | Sigma-Aldrich |
| Insulin (Humulin R) | MGH pharmacy |
| Sodium heparin | MGH pharmacy |
| Dexamethasone | Sigma-Aldrich |
| Hydrocortisone | MGH pharmacy |
| Penicillin-Streptomycin | Invitrogen |
| L-Glutathione | Sigma-Aldrich |
| Bovine Serum Albumin (BSA) | Sigma-Aldrich |
| 35kDa Polyethylene glycol (PEG) | Sigma-Aldrich |
| 3-O-methyl glucose | Chem-Impex |
| D-(+)-Trehalose dihydrate | Sigma-Aldrich |
| Glycerol | Fisher Scientific |
| Antifreeze glycopeptides (AFGP) | A/F Protein Inc. |
| X/Z-1000 | 21st Century Medicine |
| Snomax | Telemet |

